# Simian Immunodeficiency Virus and Antiretroviral Therapy Impact Rhesus Macaque Brain Lipid Distribution

**DOI:** 10.64898/2026.04.03.716347

**Authors:** Cory J. White, Kyle A. Vanderschoot, Dalton R. Brown, Alyssa F. Espley, Elizabeth K. Neumann, Caitlin M. Tressler, Dionna W. Williams

## Abstract

Human immunodeficiency virus (HIV) infection promotes considerable bioenergetic, spatially heterogenous strain to the brain that is incompletely ameliorated through viral suppression afforded by antiretroviral therapy (ART). Disrupted homeostasis of brain lipids after HIV in humans or simian immunodeficiency virus (SIV) infection in rhesus macaques occurs due to elevated energetic demands, neuroinflammation, reactive oxygen species, and barrier leakiness. Brain lipids are particularly vulnerable to HIV-associated dysregulation due to their high abundance, unique composition, and specialized functional roles. Using rhesus macaques exposed to SIV and ART (tenofovir disoproxil fumarate (TDF), emtricitabine (FTC), and dolutegravir (DTG), we investigated the spatial distribution and abundance of lipids across brain regions and metabolically relevant peripheral tissues using mass spectrometry imaging. When comparing lipid abundance, individual lipids representing a multitude of species were more varied across tissues than by treatment condition.

Further, we discerned either solely SIV infection or ART outweighed one another in altering phospholipids in different tissues Presence of ART had a greater influence on phospholipid homeostasis in the temporal cortex and hippocampus than in the midbrain, possibly due to differences in penetrance and turnover of ART across brain regions.

Overall, these data demonstrate ART robustly increased phospholipids across brain regions while SIV infection had a varied impact depending on the brain region. These findings inform the need to further evaluate the neurologic consequences that may result in the brain due to disrupted lipid homeostasis across ART regimens.

## INTRODUCTION

The human brain is a complex and energetically expensive organ that relies on precise metabolic regulation to maintain region-specific roles. Brain lipid composition is pronouncedly unique compared other tissues, with high μ-3 and μ-6 polyunsaturated long- and very long-chain fatty acids (PUFAs) that are critical for proper neurodevelopment ^1,2^. Of note, lipids are most abundant in the brain compared to all other organs, except adipose, comprising approximately 50-60% of total dry brain mass^3^. Furthermore, brain lipids are necessary for proper neuronal structure, signaling, and function ^1,2,4,5^. The essentiality of appropriate brain lipid composition is evidenced by the many neurologic diseases hallmarked by dysfunction or disruptions in lipid metabolism, including that which occurs following infectious pathogens ^6–15^.

HIV is well known to impact brain lipid composition, lipid bioenergetics, and metabolic homeostasis. While HIV enters the brain within the first two weeks of infection, these adverse lipid consequences persist for decades after initial infection^11,16^ . These disruptions in lipid metabolism are associated with neurological sequelae that persist even with viral suppression achieved by ART, suggesting long-term and persistent dysregulation of brain lipid composition and aberrant metabolic function^11^. As such, a more comprehensive understanding of lipid perturbations during HIV may be helpful in identifying potential interventions or therapeutics to restore neurologic function following infection.

It is also important to consider the contribution of ART on brain lipids, as it is recommended upon HIV diagnosis to suppress virus and used by 77% of people living with HIV^17^. Furthermore, some ART drugs are used in uninfected people as prophylaxis to prevent HIV acquisition. As these xenobiotics require lifelong adherence to treat or prevent HIV infection, it is critical to understand their metabolic and neuronal implications. Previous studies support that ARTs adversely impact bioenergetics and metabolic regulation in the brain and on peripheral tissues, though less is known about their impact on brain lipids. For example, efavirenz is a reverse transcriptase inhibitor that historically was widely used to treat HIV, but also contributes to weight gain and dyslipidemia, hepatoxicity, and elevated cholesterol^18,19^. Furthermore, in brain, efavirenz disrupts neuronal mitochondrial function, oxygen consumption, and glucose metabolism, as well as contributes to neuronal autophagy and neurotoxicity ^20–24^.

Similar findings occur with ART commonly used in contemporary regimens, including the integrase inhibitor DTG, which is associated with neuroinflammation, neural tube defects, and changes in glucose metabolism ^25–27^. However, the direct impact of ART on brain lipid metabolism, particularly with respect to the heterogenous composition of lipids in a region-specific fashion, is not well characterized.

Lipid metabolism within the brain is heterogenous and dynamic wherein differing regions have distinct spatial lipid compositions, bioenergetic needs, and regulation of metabolic homeostasis ^28–30^. Similarly, HIV infection is compartmentalized within the brain wherein virus is more frequently detected in specific brain regions. For example, basal ganglia, thalamus, and prefrontal cortex are preferentially prone to atrophy due to HIV infection ^31^. Correspondingly, ART is not always uniformly distributed in the brain.

We and others identified unequal emtricitabine, tenofovir, and DTG distributions ^32–36^. Despite the well-established variation in HIV infection and ART disposition in the brain, their individual and combined impacts on brain lipids across multiple brain regions is incompletely understood. We previously identified heterogenous distribution of membrane phospholipids and acylcarnitines across brain regions in a SIV-infected, ART-treated rhesus macaque ^30^. Now, we expand on this prior work by evaluating macaques that are 1) uninfected, 2) SIV-infected, 3) SIV-infected with viral suppression by ART, and 4) SIV-infected with treatment interruption to facilitate viral rebound. This project uncovers that PC and PE phospholipids in the brain are generally increased after any ongoing or previous exposure to ART, while SIV infection also has the capacity to alter lipid composition in the brain albeit less robustly. The changes to spatial lipid composition due to SIV and ART may result in neurologic impairments and may inform health considerations for future ART regimens.

## Materials and Methods

### Animal Ethics Statement

All procedures were performed in accordance with the *NIH’s Guide for the Care and Use for Laboratory Animals,* the *US Department of Agriculture Animal Welfare Act*, and under the approval of the Emory University Animal Care and Use Committee and Division of Animal Resources.

### SIV-infected Rhesus Macaque Model

We used a well-established SIV-infected rhesus macaque (*Macaca mulatta*) model optimal for studying brain pathology, immune compromise, viral replication, and viral suppression afforded by ART^30^. A total of four rhesus macaques were used in this study with a single macaque representing the 1) uninfected and untreated, 2) SIV only, 3) virally suppressed, and 4) viral rebound groups respectively. The present study was conducted using male rhesus macaque samples as previously described ^30,34^. All rhesus macaques were ∼2 to 3 years of age. Three rhesus macaques were inoculated intravenously using SIVmac251 as previously described^30,34^ . Two weeks after SIV infection, two rhesus macaque was treated daily with intramuscular injections of an ART regimen of tenofovir disoproxil fumarate (TDF) (20 mg/kg), dolutegravir (DTG) (2.5 mg/kg), and emtricitabine (FTC) (40 mg/kg) ^37^. ART was donated by Gilead (Foster City, CA) and ViiV Healthcare (Raleigh, NC). This model achieved peak viremia at 14 days post inoculation and accomplished complete viral suppression by day 42 post inoculation (28 days after ART initiation). At 90 days of treatment, one of the two macaque received ART treatment interruption to facilitate viral rebound, as evidenced by three consecutive positive qPCR results quantifying SIV_Gag_ RNA. All macaques were monitored to ensure there were no signs of pain or distress. The single rhesus macaque that was infected with SIV only with no ART intervention was euthanized prior to the virally suppressed and viral rebound macaques for humane reasons. Criteria for humane euthanasia prior to planned endpoint included weight loss of greater than 15%, CD4^+^ T-lymphocytes count less than 5% of baseline level, clinical signs of neurological disease, intractable diarrhea, and opportunistic infection. These criteria were not met for any animals involved in this study.

### Necropsy and Organ Collection

At approximately 130 days post inoculation, the juvenile SIV-infected, ART-treated rhesus macaque was euthanized in accordance with federal guidelines and institutional policies ^37^. Euthanasia occurred with an overdose of sodium pentobarbital while under ketamine sedation (15- to 20- mg/kg intramuscular injection) before perfusion with phosphate-buffered saline (PBS) (Gibco) to remove blood from tissues ^37–40^. The necropsy was performed per previously established protocols, in accordance with the American Veterinary Medical Association guidelines for euthanasia of animals ^37–40^.

After harvest, the whole brain was placed in cold 2.5% agarose, and 4mm coronal sections were obtained and annotated in accordance with the rhesus macaque Scalable Brain Atlas ^41^. Brain regions (frontal cortex, temporal cortex, thalamus, cerebellum and midbrain) and peripheral tissues (kidney, liver, and spleen) were fresh frozen using liquid nitrogen in optimum cutting temperature medium (OCT) (Sakura Finetek Inc, Torrance, CA) and stored at -80°C until cryosectioning ^38^.

### Tissue Sectioning and Matrix Application

The MALDI matrix, a-cyano-4-hydroxycinnamic acid (CHCA), was obtained from MilliporeSigma (St. Louis, MO). The CHCA matrix is widely suitable for lipids ^42^. All solvents and other chemicals used were either reagent or high-performance liquid chromatography (HPLC) grade, and purchased from Fisher Scientific (Hampton, NH), unless otherwise specified.

OCT-embedded cerebellum, frontal cortex, hippocampus, kidney, liver, midbrain, spleen, and temporal cortex were sectioned to 10 μm onto MALDI Intellislides (Bruker Daltonics, Bremen, Germany) at -20°C using a cryostat (Leica Biosystems, Buffalo Grove, IL). Matrix [CHCA, 10 mg/ml in ACN:H_2_O (50:50, v/v)] with 0.03% (v/v) TFA was applied to each slide using a HTX M5 Sprayer (HTX Technologies, LLC, Chapel Hill, NC) at a flow rate of 100 mL/min. The TM-Sprayer was operated at an air pressure (N_2_) of 10 psi, spray nozzle velocity of 1200 mm/min, track spacing at 2 mm, and spray nozzle temperature at 80°C. For homogeneous deposition of matrix onto each slide, we used 16 passes (matrix deposition cycles). One 10-um coronal slice of tissue from each representative macaque group was placed on an individual slide, when possible, to limit batch effects.

### MALDI-IMS Data Acquisition

MALDI-IMS data was acquired in the Applied Imaging Mass Spectrometry Core/Service Center at Johns Hopkins University School of Medicine using the samples on MALDI Intellisides. All slices to be compared were prepared on a single slide for simultaneous acquisition at 50-micron resolution using a Bruker Rapiflex MALDI TOF/TOF instrument in positive mode.

### Selection of Ions and Analysis

Spectra data was acquired and normalized by total ion count in SCiLS (Bruker Scientific). Heat map using manual traces from brain and peripheral tissue slices were generated using SCiLS software. For each species of every tissue, images were captured for H^+^, Na^+^, and K^+^ adducts. Receiver operating characteristics (ROC) plots between all groups were used to determine lipids of interest. Peak area for each lipid was used for statistical analysis. LipidMAPS database searching was used to identify lipids associated with ions of interest.

### Tandem Mass Spectrometry

Additional tandem mass spectrometry (MS/MS) in positive mode and on-tissue fragmentation was used to validate the identify of several lipids from MALDI-IMS tissue slides (Figure S2). Annotation of lipids were performed manually through identification of parent lipid and fragment matches using LipidMaps.

## Results

Brain lipid composition is most frequently evaluated using homogenized tissue. However, this approach cannot provide insight regarding the spatial distribution of lipids within intact brain microstructure. To evaluate the impact SIV infection and ART treatment on spatial arrangement of lipids in the brain, we used MALDI-IMS to map the spatial distribution of ions across brain regions prone to higher viral load and atrophy following HIV infection (hippocampus, temporal cortex, midbrain, frontal cortex, cerebellum)^34,46^. Importantly, the ions mapped were specifically those whose masses fall within the typical mass range and are indicative of a wide array of common lipids (100 Da – 1000 Da; glycerophospholipids, sphingolipids, and fatty acid derivatives). As the brain has a unique lipid composition compared to other tissues, we also evaluated HIV-and metabolically relevant peripheral tissues (liver, kidney, spleen) in the same manner to evaluate whether there were changes in lipid composition that were brain-specific or that occurred more broadly across organs. For this study, we leveraged the well-established rhesus macaque model that included four treatment groups: 1) uninfected, 2) SIV-infected, 3) SIV-infected with viral suppression by ART and 4) SIV-infected with treatment interruption to facilitate viral rebound. Coronal slices of tissue from each representative macaque group were placed on an individual slide. Slides containing tissue slices from each treatment group were prepared and MALDI-IMS performed with a 50μm resolution (**Figure 1A**), acquiring ions at a mass range (100Da to 1000Da) corresponding to common lipids: glycerophospholipids (PCs, PEs, PSs, PGs, and PAs), sphingolipids (sphingomyelins and ceramides), and CARs. As disrupting the metabolic homeostasis of these classes of lipids in the brain has been associated with neurologic consequences^47–49^, it is important to assess the effect of HIV and ART administration on lipid abundance and distribution in the brain.

**Figure 1.**
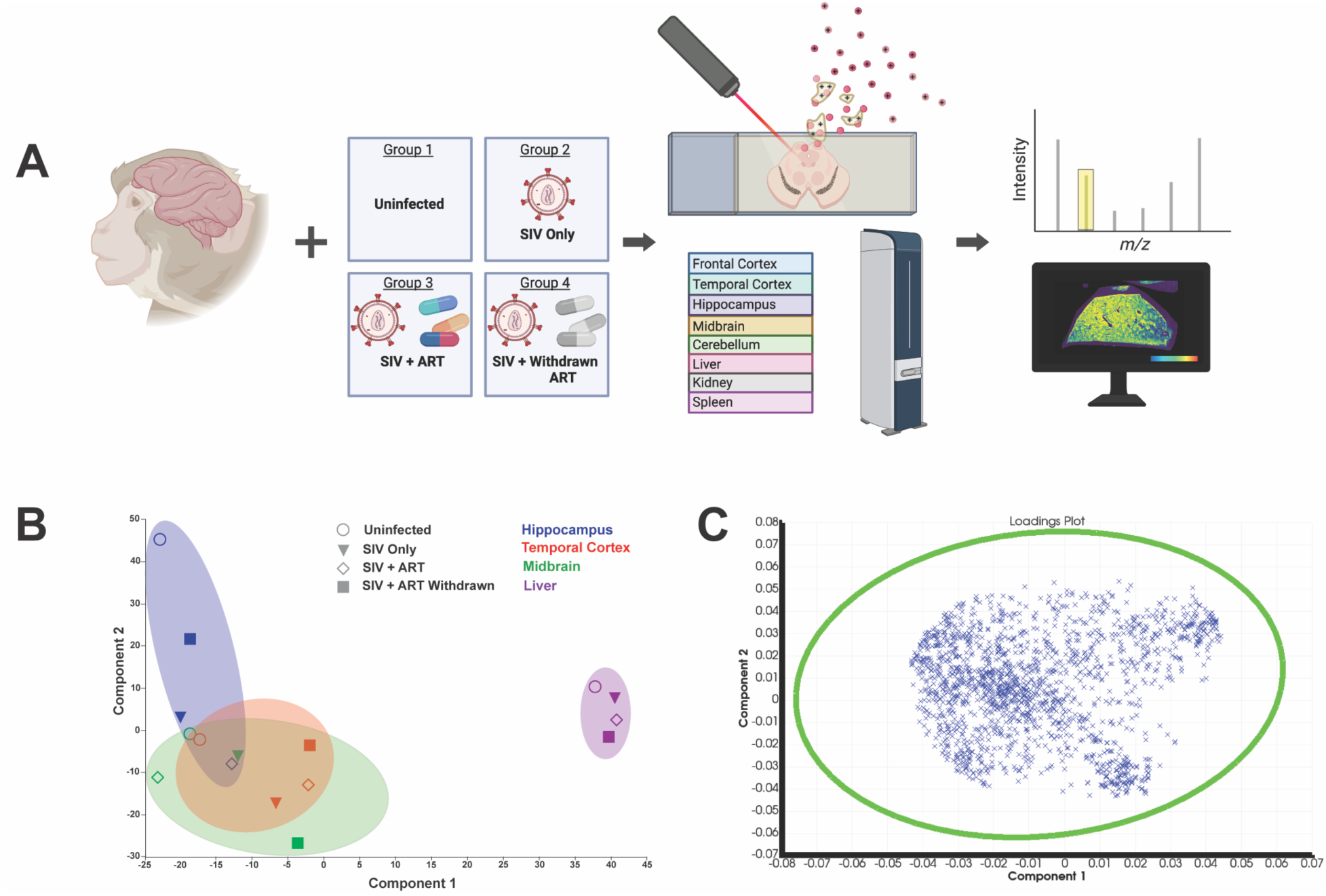
Spatially Mapping the Brain of SIV Rhesus Macaque Model Across Conditions of Infection and Treatment. (A) Schematic representing the four rhesus macaque models representing four permutations of infection status and ART treatment including Group 1: Uninfected (uninfected, untreated control), Group 2: SIV Only (SIV infected, untreated), Group 3: SIV + ART (*virally suppressed* SIV-infected, ART-treated), and Group 4: SIV + Withdrawn ART (*viral rebound* SIV-infected, formerly ART-treated). From each representative group, if available, brain regions (frontal cortex, temporal cortex, hippocampus, midbrain, and cerebellum) and peripheral tissues (liver, kidney, and spleen) were imaged using a Bruker Rapiflex with CHCA matrix and positive mode. The representative chromatogram illustrates how a spatial intensity map can be generated from a singular *m/z* value. In the representative spatial intensity image, red signifies the highest intensity, and blue signifies the lowest of each *m/z* value. (B) Principal component analysis (PCA) of the four tissues [hippocampus (blue), temporal cortex (red), midbrain (green), and liver (purple)] with complete representation across all groups [Uninfected (open circle), SIV Only (shaded triangle), SIV + ART (open diamond), SIV + ART Withdrawn (shaded square)]. (C) Example loadings plot for total data.

### Lipid Composition in Brain Tissues Remains Distinct to Peripheral Tissues Regardless of SIV Infection or ART Treatment

In our first set of analyses, we performed principal component analysis (PCA) using the total acquired ion data to identify variance and similar trends in lipid distribution across all tissues and treatment groups. When comparing the PCA of total ions across tissues, we determined that the hippocampus had a distinct lipid ion profile compared to other brain regions, regardless of treatment group (**Figure 1B**). As expected, we determined that distinct ion profiles occurred between brain (hippocampus, temporal cortex, and midbrain) and liver confirming that the brain possesses unique lipid composition relative to peripheral organs (**Figure 1B**).

Additionally, variance between organs was generally greater than between treatment groups for hippocampus and liver (**Figure 1B**). When assessing loading plots from all tissues to determine the contributions of individual ions to overall variance in loading plots we observed no distinct ions, indicating no overwhelming influence by individual ions to overall variance across all the acquired data (**Figure 1C**).

### Lipid Species are Uniquely Affected by SIV Infection and ART

We next sought to disentangle the effect of SIV infection and ART treatment on global lipid species representing phospholipids, sphingolipids, and CARs in brain.

Firstly, using peak area normalized to total ion count, we assessed variance in the abundance of individual lipid species across all treatment groups amongst all tissues. Multiple lipid species were elevated across brain regions including CAR: CAR22:3(10Z,13Z,16Z); sphingolipid: SM(d34:1)+H and CerP(36:2; O_2_)+K, and phospholipids PC(P-34:2)+H and PC(36:3)+K (Figure S1). CAR22:3(10Z,13Z,16Z) and CerP(36:2; O_2_)+K, trended higher, albeit not significantly, in hippocampus than in midbrain and temporal cortex (Figure S1). Other lipid species were not robustly altered but exhibited distinct patterns amongst brain and peripheral tissue. For example, phospholipid PC(36:2)+Na was significantly higher in hippocampus compared to temporal cortex and its abundance in spleen reached levels more like brain than other peripheral organs (Figure S1). Uniquely, PC(36:2)+Na was also significantly elevated in kidney compared to temporal cortex and CAR4:0 was comparably abundant in spleen, hippocampus, midbrain, and temporal cortex (Figure S1). Both paradigms indicate lipid species predominant in brain can also be found at comparable levels in singular peripheral tissues. Other lipid species were preeminent in liver compared to all other tissues including phospholipids PC(34:2)+Na, PC(34:0)+Na, PC(36:1)+Na, PA(36:2)+K, PC(34:1)+Na, and PC(32:0)+K as well as sphingolipids SM(d42:2)+H (Figure S1).

Similarly, CAR2:0 predominated in kidney, spleen, and liver while lipoxygenation metabolite 15-HETrE predominated in kidney and spleen (Figure S1). Overall, these data suggest that individual lipid species play tissue specific roles to maintain proper function.

We next looked across treatment groups using data from the above analyses to investigate if SIV infection or ART impacted the presence of individual lipid species in each tissue. Across these data, we observed trends but no consistent pattern to a class of lipids in any tissue or treatment group. For example, the following lipid species were highest in virally suppressed hippocampus including SM(38:1), PC(32:0)+Na, PC(36:2)+H, PE(38:1)+H PC(36:3)+K, PE(38:2)+H, CerP(36:2; O_2_)+K, 15-HETrE, PC(34:2)+Na, PC(32:0)+H (Figure S1). Further in hippocampus, several adducts representing lipid species in the viral rebound group, including phospholipids PC(32:0)+Na, PC(32:0)+H, PC(36:3)+K, PC(36:2)+H, PE(38:1)+H, PE(38:2)+H; sphingolipid, CerP(36:2; O_2_)+K; and lipoxygenation metabolite 15-HETrE, were decreased compared to other treatment conditions (Figure S1). This may indicate that ART is not only necessary to restore these lipids to baseline but increases them beyond endogenous levels.

Incongruent trends were observed in other brain regions. In midbrain, lipid species were often highest in the SIV-only group including PC(36:3)+K, PC(32:0)+Na, PC(36:2)+H, PC(34:0)+Na, PC(32:0)+K, PC(P-34:2)+H CAR22:3(10Z,13Z,16Z), and SM(d42:2)+H (Figure S1). However, there was no consistent treatment group with a lowest lipid intensity. These elevated phospholipids in the SIV only group could be indicative of incomplete ART penetrance, as the midbrain is in deeper brain tissue. Collectively, these data may indicate that the individual lipid species may be regulated distinctly by HIV infection and ART regimen, or lack thereof.

The following paragraphs are further representative demonstrations of how lipid is differentially regulated by SIV and ART

### Lipids are Distinctly Regulated by SIV or ART Across Brain Regions and Liver

To investigate further, normalized receiver operating curves (ROCs) were used to determine individual lipids that discriminate due to SIV infection and ART treatment in brain regions and liver. Firstly, in the temporal cortex, we identified 18-oxo-9S,15S-dihydroxy-11R-acetoxy-5Z,13E prostadienoic acid as a lipid reflective of SIV infection as it was decreased in the setting of infection but restored with viral suppression.

Interestingly, viral rebound following treatment interruption did not restore the lipids to endogenous levels, suggesting prolonged viral replication (rather than short bursts of viral production) may be required to decrease this lipid (**Figure 2A**). In contrast, PC(36:2) was most prominent in any animals treated with ART regardless of if therapy was stopped or viral rebound occurred, suggesting a long-lasting effect on regulation of this lipid (**Figure 2B**). A similar yet inverse trend occurred in liver, where ART had a pronounced effect in decreasing SM(d42:2)H+ that was unable to be restored after stopping ART (**Figure 2E**). Additionally, we determined SM(42:1;O_2_) Na^+^ was increased only in hippocampus that were both SIV-infected and treated with ART, indicating a distinct pattern at this intersection (**Figure 2D**). Importantly, some lipids were incredibly stable and resilient to modulation by either infection or ART treatment, including SM (d34:1) in the midbrain (**Figure 2C**). These findings further demonstrate a heterogenous impact of SIV infection and ART treatment on the lipid composition of individual lipid species. Where some are resistant to modulation, others are shifted in response to viral replication, while still other species were sensitive specifically to ART.

**Figure 2.**
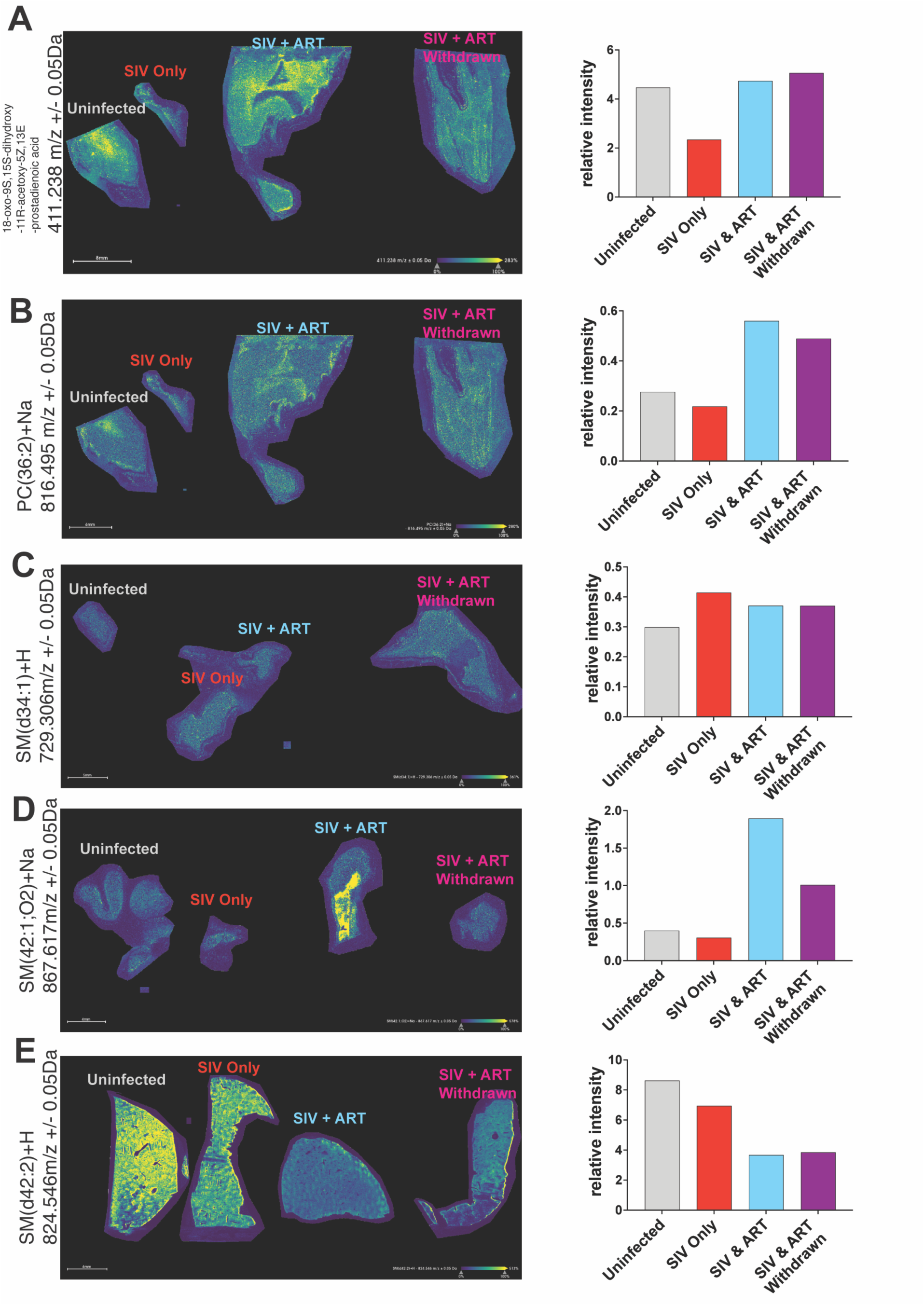
Abundance for Individual Lipid Species in Multiple Brain Regions Follow Trends by Infection Status and Antiretroviral Regimen. Representative MALDI-IMS of representing the four rhesus macaque models representing four permutations of infection status and ART treatment including Group 1: Uninfected (uninfected, untreated control), Group 2: SIV Only (SIV infected, untreated), Group 3: SIV + ART (*virally suppressed* SIV-infected, ART-treated), and Group 4: SIV + Withdrawn ART (*viral rebound* SIV-infected, formerly ART-treated). (A & B) are from temporal cortex, (C) is from midbrain, (D) is from hippocampus, and (E) is from liver. Red signifies the highest intensity, and blue signifies the lowest of each *m/z* value.

### ART Does Not Restore SIV-Induced Perturbations to Spatial Lipid Abundance in the Midbrain

We next assessed the impact of SIV infection and ART treatment on the intensities of individual lipids in the midbrain, a brain region that is disproportionately infected by HIV and prone to atrophy in humans, using ROCs^34^. Several increased lipids, including ions PC(26:0)+H^+^, PE(38:2)+H^+^, SM(d34:1)+Na^+^, and SM(d34:1)+H^+^, were indicative of unique factors contributing to increases in the SIV-infected and virally suppressed groups compared to the uninfected and viral rebound groups. The decrease in the viral rebound group was particularly unexpected, as both infection and any ART exposure were increased abundance of these lipids in the SIV-infected and virally suppressed groups (**Figure 3**). This data demonstrates that in midbrain, SIV potentially has a greater influence on altering lipid than ART.

**Figure 3.**
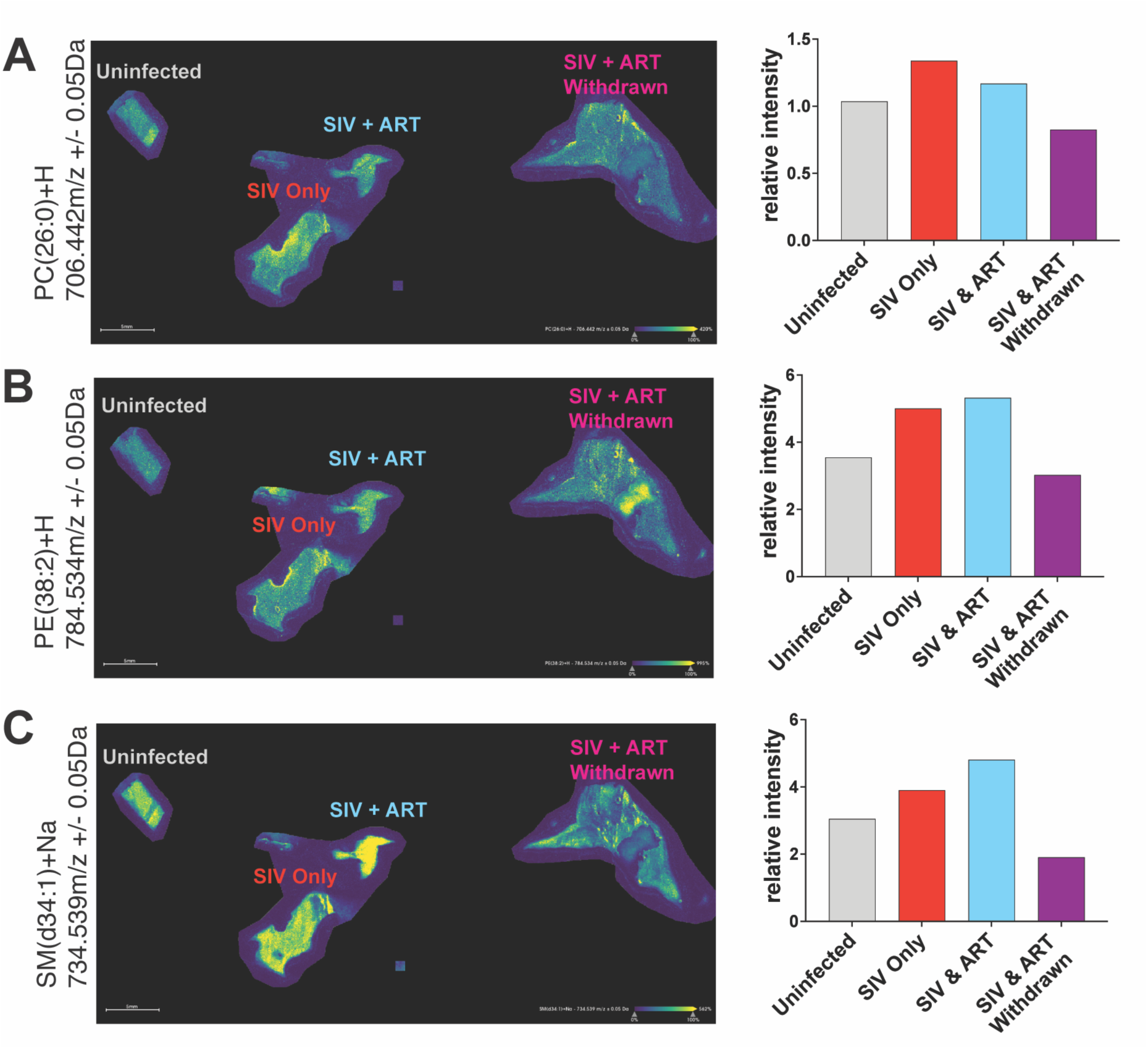
Spatial Lipid Abundance is Elevated in SIV Only and SIV+ART Rhesus Macaque Midbrain. Representative MALDI-IMS of representing the four rhesus macaque models representing four permutations of infection status and ART treatment including Group 1: Uninfected (uninfected, untreated control), Group 2: SIV Only (SIV infected, untreated), Group 3: SIV + ART (*virally suppressed* SIV-infected, ART-treated), and Group 4: SIV + Withdrawn ART (*viral rebound* SIV-infected, formerly ART-treated). (A-C) are from midbrain. Red signifies the highest intensity, and blue signifies the lowest of each *m/z* value.

### SIV Infection Increases Phospholipids in Temporal Cortex, while ART Depletes Distinct Liver Phospholipids

Comparing ROC data across temporal cortex and hippocampus, we investigated the impact SIV infection and ART treatment on phospholipids, which are a major component of cellular and organelle membranes, across brain regions and liver that were tissues with representation across all groups. In temporal cortex, lipids that were good discriminators were increased in the SIV-infected only groups including PG(O-32:0)+K^+^, PE(36:1)+H^+^, PC(34:1)+H^+^, PC(P-34:1)+Na^+^ were more abundant in the SIV-infected group in temporal cortex (**Figure 4A**). Additionally in hippocampus, the lipids PC(36:2)+Na+ and PC(34:1)+H+ were determined by ROC to be elevated.

**Figure 4.**
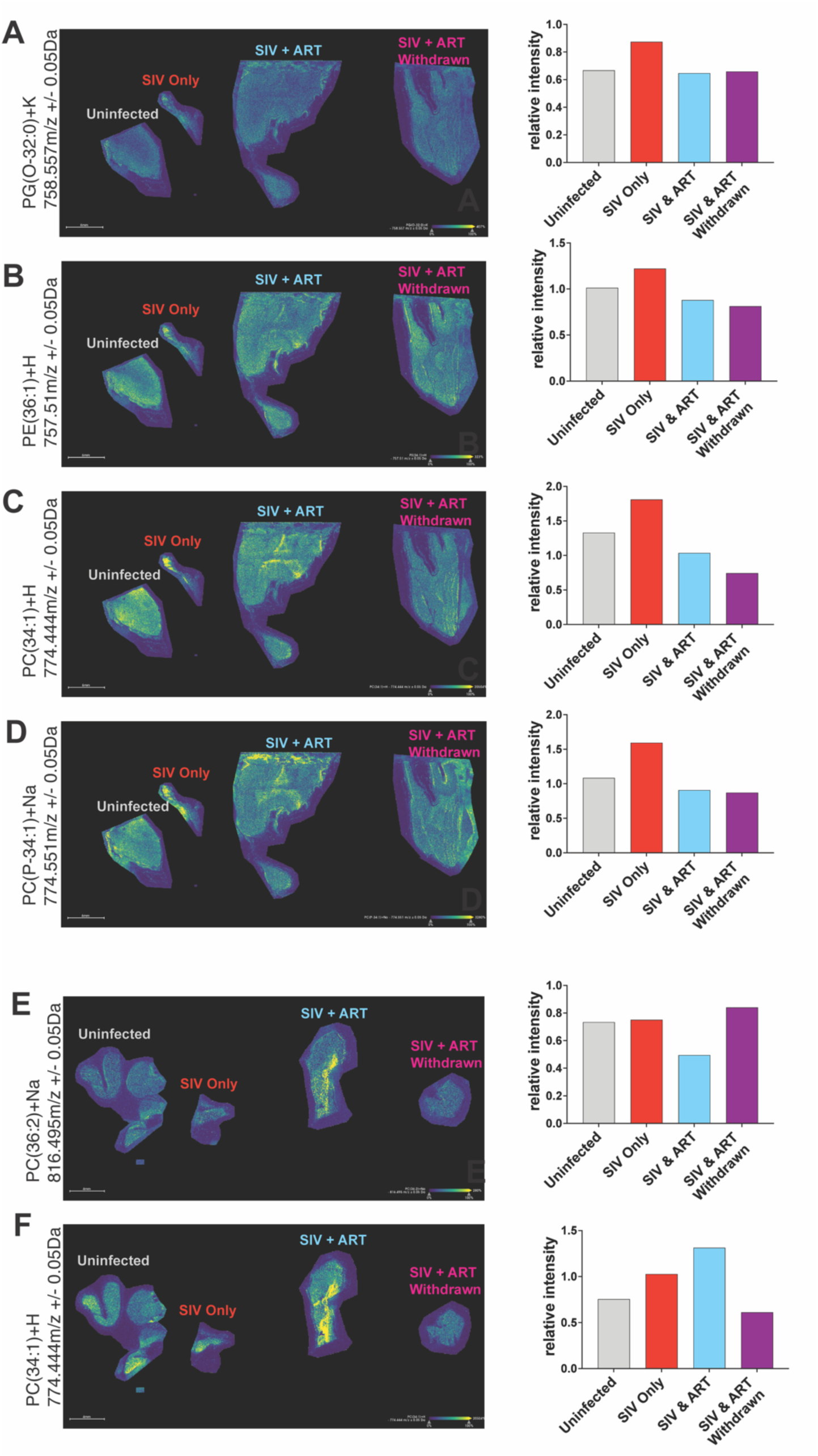
Temporal Cortex Phospholipids Are Increased by SIV Infection. Representative MALDI-IMS of representing the four rhesus macaque models representing four permutations of infection status and ART treatment including Group 1: Uninfected (uninfected, untreated control), Group 2: SIV Only (SIV infected, untreated), Group 3: SIV + ART (*virally suppressed* SIV-infected, ART-treated), and Group 4: SIV + Withdrawn ART (*viral rebound* SIV-infected, formerly ART-treated). (A-D) are from temporal cortex. (E-F) are from hippocampus. Red signifies the highest intensity, and blue signifies the lowest of each *m/z* value.

PC(36:2)+Na+ was decreased solely in the virally suppressed group (**Figure 4B**). Interestingly, PC(34:1)+H was more elevated in the virally suppressed group in hippocampus. Further, we assessed SIV infection and ART treatment in liver. Conversely to brain regions, most lipids in liver, found distinct due to SIV infection and ART treatment by ROC, were higher in groups that had no history of ART treatment. For example, PC(34:1)+H+, PC(34:1)+Na+, SM(d34:1)+K+, and PC(36:1)+Na+ were all higher in the uninfected group and SIV-infected only group (**Figure 5**). However, deviating from this pattern, PC(32:2)+H+ was most abundant in the virally suppressed group (**Figure 5**). Together, these data displayed that lipids in temporal cortex were increased by SIV infection, liver was decreased by ongoing or previous ART, and lipids in hippocampus had no particular trend (**Figure 4B**; **Figure 5**). This further demonstrates that individual lipids are modulated distinctly by SIV infection and ART treatment across both the brain and peripheral tissues.

**Figure 5.**
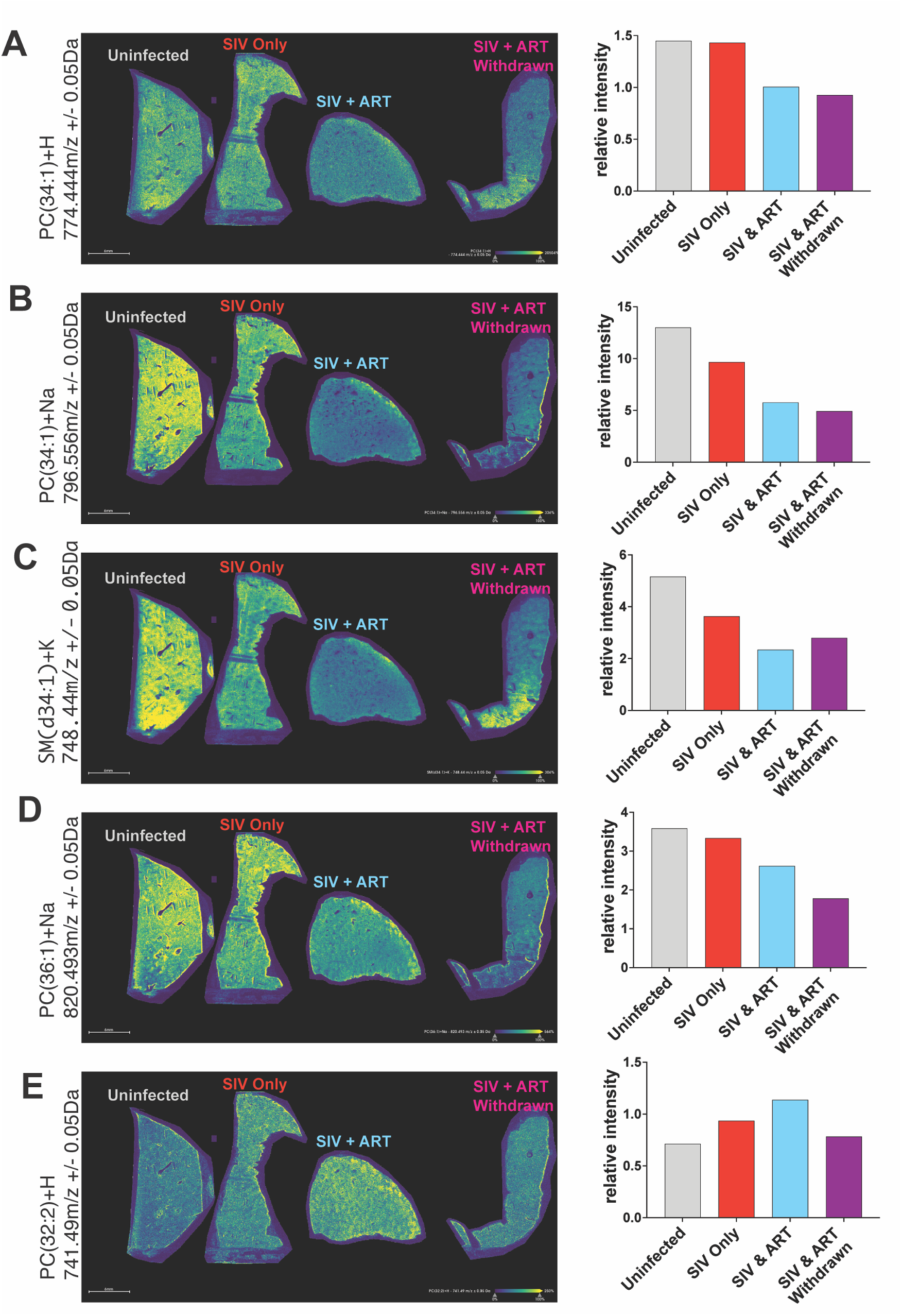
No Specific Pattern in Spatial Lipid Composition is Observed in Liver. Representative MALDI-IMS of representing the four rhesus macaque models representing four permutations of infection status and ART treatment including Group 1: Uninfected (uninfected, untreated control), Group 2: SIV Only (SIV infected, untreated), Group 3: SIV + ART (*virally suppressed* SIV-infected, ART-treated), and Group 4: SIV + Withdrawn ART (*viral rebound* SIV-infected, formerly ART-treated). (A-E) are from liver. Red signifies the highest intensity, and blue signifies the lowest of each *m/z* value.

### PC and PE are Generally Increased by ART and SIV in Midbrain but Decreased by SIV in the Hippocampus and Temporal Cortex

We determined that individual lipids are regulated by SIV infection and ART treatment across brain regions and peripheral tissues. However, trends across common lipid species such as phospholipids, that comprise cellular membranes, may inform molecular consequences relevant to neuropathology. To comprehensively evaluate global changes in brain phospholipids among all treatment groups, we generated a graphical representation of species from the most abundant classes of phospholipids, PCs and PEs in hippocampus, midbrain, and temporal cortex by creating a rendering based on the above-described ROC analyses^50^. First, we assessed the impact of SIV infection on PC and PEs. We determined that SIV infection decreased PC and PE abundance in hippocampus and temporal cortex, relative to uninfected macaques. However, in midbrain PC and PE abundance was elevated in SIV-Infected Only (**Figure 6A**).

**Figure 6.**
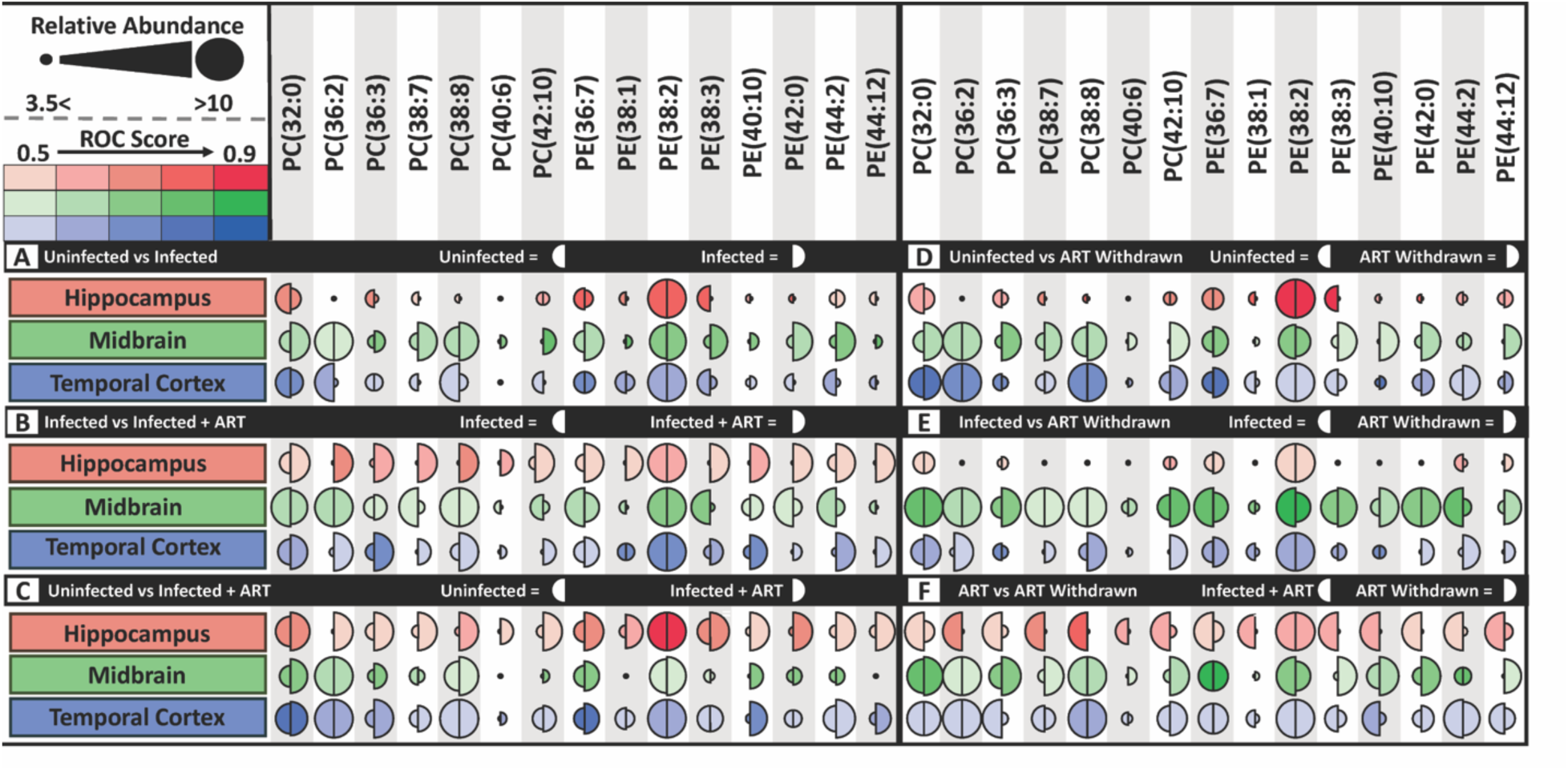
Ongoing and Previous ART Exposure Both Increase PCs and PEs in Hippocampus and Temporal Cortex. This bubble plot describes changes in lipid composition across conditions of SIV-infection and ART treatment in hippocampus (red), midbrain (green), and temporal cortex (blue). Lipid species are described in each functional unit by two half circles correlating their relative intensity and ROC value. The larger the half circle, the more abundant the lipid species in that treatment condition. Darker shading indicates a great ROC value. (A) Uninfected (left half circle) vs. SIV-infected (right half circle) (B) SIV-infected (left half circle) vs. SIV-infected + ART-treated (right half circle) (C) Uninfected (left half circle) vs. SIV-infected + ART-treatment (right half circle). (D) Uninfected (left half circle) vs SIV-infected + ART-treatment Withdrawn, E) SIV-infected (left half circle) vs SIV-infected + ART-treatment Withdrawn, (F) SIV-Infected + ART, SIV-infected + ART Withdrawn.

The influence of ART was assessed when SIV-infected Only and virally suppressed groups were compared. We found that virally suppressed PEs and PCs from the hippocampus and temporal cortex were elevated compared to SIV-infected Only groups (**Figure 6B**). Inversely, making this sample comparison in midbrain, PC and PEs in SIV-infected only samples were elevated (**Figure 6B**). The combined influence of SIV infection and ART treatment in PCs and PEs was assessed by comparing the virally suppressed group to the uninfected group. Both together result in elevated PCs and PEs in hippocampus and temporal cortex (**Figure 6C**). This finding mirrors the impact of ART alone found in hippocampus and temporal cortex (**Figure 6B**). However, in virally suppressed compared to uninfected midbrain the influence of SIV infection is observed as infection increases PCs and PEs (**Figure 6C**). This finding is similar to how midbrain was impacted by SIV infection only (**Figure 6A**). Further, similar trends were observed to Figure 5B when comparing SIV Only to virally suppressed brain regions (**Figure 6C**). Consistency between these brain regions suggest they are impacted by SIV infection and ART treatment in similarly. Next, we sought to assess the impact of viral rebound and a history of previous ART exposure. When comparing the uninfected group, which was ART naïve, to viral rebound group, which had a previous ART exposure, previously congruencies in trends across tissues were not upheld. Viral rebound (or previous ART exposure history) decreased PC and PE species in hippocampus (**Figure 6D**). However, in both midbrain and temporal cortex, PC and PE species were increased in the viral rebound group (**Figure 6D**). We next assessed PCs and PEs in ongoing SIV infection compared to viral rebound. In midbrain or hippocampus ongoing SIV infection or viral rebound, which had a previous ART history had a minor impact overall on PC and PE abundance (**Figure 6E**). This suggests that something unique to SIV infection may be impacting phospholipid regulation in these brain regions. However, in temporal cortex, PC and PE species in the viral rebound group were generally more abundant than SIV-infected only group (**Figure 6E**). This supports that ART had more influence on lipid regulation in temporal cortex. Lastly, the impact of ongoing ART treatment vs previous ART exposure as well as viral suppression on PC and PE regulation was assessed by comparing virally suppressed and viral rebound hippocampus, midbrain and temporal cortex. In hippocampus, virally suppressed PC and PEs are pronouncedly more abundant than those in the viral rebound group (**Figure 6F**). This could suggest that in hippocampus either SIV infection decreased these PCs and PEs or that ARTs, which were actively being administered in the virally suppressed group, increased PCs and PEs. Similarly, in temporal cortex, virally suppressed PCs and PEs are more abundant, albeit less so than hippocampus, than viral rebound (**Figure 6F**). Together, these data demonstrate that ART treatment has an outsized impact of phospholipid abundance as groups with ART exposure generally are increased.

### Patterns in PC and PE species in kidney, liver, and spleen observed across treatments are unique in each organ

We evaluated the impact of SIV infection and ART treatment across PC and PE species in the peripheral tissues kidney, liver, and spleen, which are all metabolically relevant and subject to injury following HIV infection, as a comparison to the aforementioned brain tissue graphical representation of ROC data^50^ (**Figure 7**). Firstly, the sole contribution of SIV infection to PCs and PEs was assessed by comparing the uninfected and SIV-infected only groups. In kidney, PCs and PEs remained unchanged due to SIV infection (**Figure 7A**). Using the same comparison in liver, PC species were increased in the SIV-infected only group potentially indicating PC-specific regulation in liver compared to PEs (**Figure 7A**). Next, we assessed the contributions of ART treatment to regulation of PCs and PEs in peripheral tissues by comparing the SIV-infected vs virally suppressed groups. In kidney, both PCs and PEs remained largely unchanged **Figure 7B**). However, making this comparison in liver, most of both PCs and PEs were more abundant in the SIV-Infected Only group (**Figure 7B**). Interestingly, spleen displayed the opposite trend occurred with increased PCs and PEs in the virally suppressed group (**Figure 7B**). Subsequently, we evaluated the impact of viral suppression of SIV infection with ART by comparing the virally suppressed to the uninfected group. In kidney, several PCs and PEs were decreased due to viral suppression (**Figure 7C**). Changes in liver due to viral suppression compared the uninfected group were like kidney but more robust with both PCs and PEs decreased due to viral suppression (**Figure 7C**). This suggests that unlike most other organs exposed to ART, lipid abundance decreases in liver and kidney with ART. Conversely, spleen PCs and PEs increased due to viral suppression (**Figure 7C**). Overall, while there was no consistent trend due to SIV infection amongst these peripheral organs, ART treatment decreased PCs and PEs in kidney and liver and increased these phospholipids in spleen.

**Figure 7.**
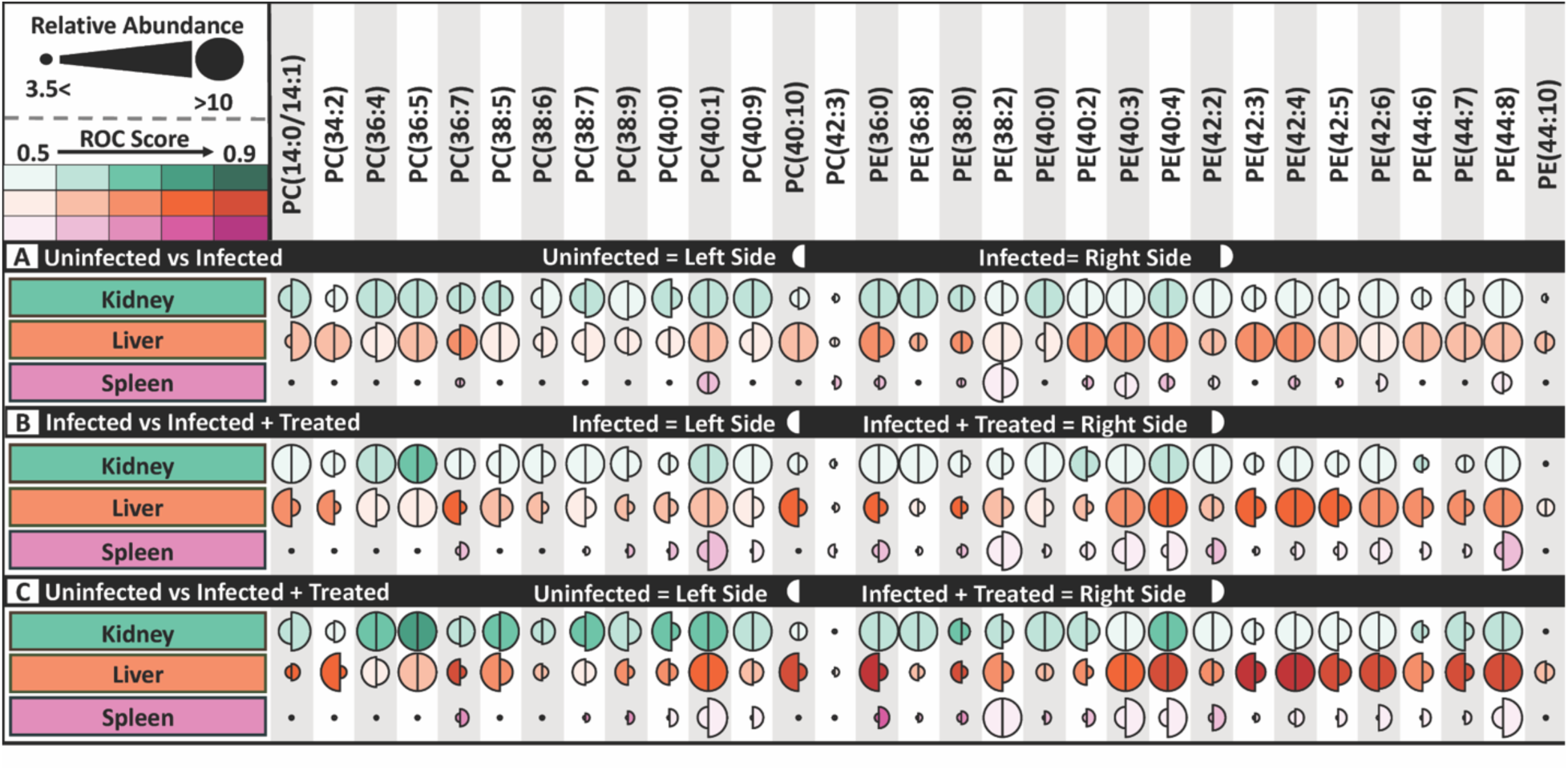
Patterns in PC and PE Species in Kidney, Liver, and Spleen Observed Across Infection and Treatment Groups are Unique in Each Organ. This bubble plot describes changes in lipid composition across conditions of SIV-infection in kidney (teal), liver (orange), and spleen (pink). Lipid species are described in each functional unit by two half circles correlating their relative intensity and ROC value. The larger the half circle, the more abundant the lipid species in that treatment condition. Darker shading indicates a great ROC value. A) Uninfected (left half circle) vs. SIV-infected (right half circle) B) SIV-infected (left half circle) vs. SIV-infected with ART-treated (right half circle) C) Uninfected (left half circle) vs. SIV-infected with ART-treatment (right half circle).

## Discussion

Brain lipids are essential for appropriate neurologic function. However, brain lipid metabolism can be disrupted by disease and by xenobiotics in a region-specific manner contributing to brain atrophy, elevated neuroinflammation, increased reactive oxygen species, energetic imbalances, and cognitive decline^3,11,31,48,49^. The impact of SIV infection and ART treatment on the spatial localization of brain lipids remain under characterized, the understanding of which can provide mechanistic insight into pathways underlying neurologic disease in people living with HIV. Our study aimed to bridge this gap by evaluating the spatial composition of lipids in brain and peripheral organs to evaluate the independent and combined effects of SIV infection and ART treatment. We found that an ART regimen of TDF, DTG, and FTC, when used currently or if treatment was stopped, contributed to long-lasting changes in phospholipid composition across multiple brain regions (hippocampus, midbrain, temporal cortex) regardless of infection status in rhesus macaques. Most notably, our study highlights that ART treatment regulates brain phospholipid metabolism more robustly than SIV infection. This was exemplified by increased PCs and PEs, prominently in hippocampus and temporal cortex, in the virally suppressed group and the viral rebound group when compared to either the uninfected or the SIV-infected only groups. Interestingly, this above trend was only observed in spleen amongst assessed peripheral tissues.

In this study we determined individual and synergistic contributions of HIV/SIV infection and ART on the spatial brain lipidome that may contribute to longstanding changes neurologic consequences^33^ Our finding that ongoing and previous ART use irreversibly alter the composition of lipids in a region-specific manner in the brain was particularly striking. The capacity for ARTs to alter brain lipid metabolic homeostasis is largely under characterized. Previously, our laboratory determined a wide breadth of phospholipids and acylcarnitines are present across brain regions in a SIV-infected, ART-treated rhesus macaque^30^. However, the influence of ART to brain lipids requires ART to cross the blood-brain barrier into the brain tissue. Evidence for spatial deposition and action of ART in the brain was demonstrated in rats after intraperitoneal injection of individual single dose administrations of 10% tenofovir, emtricitabine, or efavirenz were detectable across brain regions using MALDI-MSI^32^.

The varied regulation of lipid species across brain regions and in peripheral tissues we observed in this study may also be attributed to baseline differences and modulation of local drug metabolism of ARTs. Relevantly, our laboratory determined that the regiment of TDF, FTC, and DTG resulted in significant changes in the expression of enzymes in cultured human astrocytes and pericytes of several pathways such as nucleotide metabolizing enzymes, potentially responsible for clearing FTC and TDF, as well as lipid metabolizing enzymes in astrocytes^34^. Different rates of clearance of ARTs by these enzymes may impact their disposition in brain and other tissues as well as affect their capacity to regulate lipid metabolism.

Additionally, in this study we see that individual lipid species display unique distribution patterns within brain regions which altogether are generally increased in brain due to ART and SIV infection. Our study is the first to assess the spatial profile of lipids in the brains of a macaque model of treated with ART ^30^. Other groups have determined that mouse tissue treated with efavirenz has altered phospholipid and sphingomyelin distribution and abundance using MALDI-MSI, primarily increasing lipid abundance^50^. While Phulara et. al focuses on efavirenz a different ART, the overall increase in phospholipid abundance is like the findings of our study ^50^.

While our study uncovered ARTs have a longstanding impact on phospholipid metabolism, most of the historic and modern literature on the impact of ART on lipids focused on peripheral tissues. Both historic and modern ART, such as the INSTI DTG, are known to contribute to weight gain which is at least in part believed to be due to altered lipid metabolism. For example, in visceral fat tissue from mice treated with tenofovir alafenamide TAF, FTC, and DTG, expression of fatty acid metabolism regulating genes Ppar-γ and leptin were significantly increased^51^. In human plasma, ART has been determined to be critical for maintaining circulating PUFAs, lysophosphatidylcholine lysophosphatidylethanolamine, and lysophosphatidic acid.

However, total elevated SFAs are increased in plasma from people treated with ART with HIV compared to uninfected or those with HIV prior to treatment^52,53^. Relevant to peripheral tissues, evidence from our study provides further support by demonstrating that ART increased the abundance of PCs and PEs in ART-treated, SIV-infected rhesus macaque kidney and spleen while decreasing PCs and PEs in liver.

Furthermore, no other studies that have compared the spatial impact of HIV/SIV infection on the brain lipidome. However, data has assessed the impact of HIV/SIV on lipids using *ex vivo* homogenized tissues from postmortem human brain and rodent models of HIV infection. In postmortem human brain, sphingomyelin and ceramides with several acyl chains C18, C20, C22, C24 were increased in HIV-infected frontal cortex, parietal cortex, cerebellum, and cerebrospinal fluid^54^. In HIV transgenic rats, PUFAs are increased in brain phospholipids, NEFAs, and TAGs^55^. Subsequent studies using our macaque models may additionally quantify changes to lipids using targeted methods to investigate substrate flux and underlying molecular mechanisms that result in altered lipid composition.

While our study uncovers a prominent effect on the abundance of brain regions across the brain due to ART treatment and less robustly SIV infection, the impact of infection and ART on brain lipid metabolism remains incomplete. Even though our study integrates spatial imaging to investigate changes in brain lipid distribution, slices could not comprehensively span across the entire brain due to tissue availability and technical constraints. MALDI-IMS could best compare regions whether within an individual brain or between animals of different treatment groups if entire brain slices from macaques representing each treatment group be imaged in the same run of the mass spectrometer. Additionally, these data also are only capable of providing a snapshot of the metabolic landscape in each tissue at a given time. Further study of substrate utilization and integration, temporal measures of metabolic consumption, and expression changes to transcription factors and enzymes that regulate metabolic homeostasis would be necessary to more understand brain lipid metabolism in the context of SIV infection and ART treatment.

## Author Contributions

C.J.W. and D.W.W. contributed to initial conceptualization and design of this study.

C.J.W. K.A.V., D.R.B., A.F.E., C.M.T., and E.K.N. performed experiments, K.A.V., D.R.B., C.M.T., and E.K.N. and developed methodologies, C.J.W. and D.W.W. wrote the original manuscript draft with input and edits from all authors.

## Acknowledgements

The authors are grateful for ample scholarly discussion with current and past members of the Williams laboratory. Research reported in this publication was supported by the National Institutes of Health under award number R00 DA044838, R01 DA052859, and U01 DA058527 (DWW), and K00 NS118713 (CJW). This work was supported, in part, by pilot funding provided by parent funding under the JHU NIMH Center for Novel Therapeutics for HIV-associated Cognitive Disorders P30 MH075673 to Justin C. McArthur. The content is solely the responsibility of the authors and does not necessarily represent the official views of the National Institutes of Health. Graphical images in figures were created using BioRender.

---

**Figure S1.**
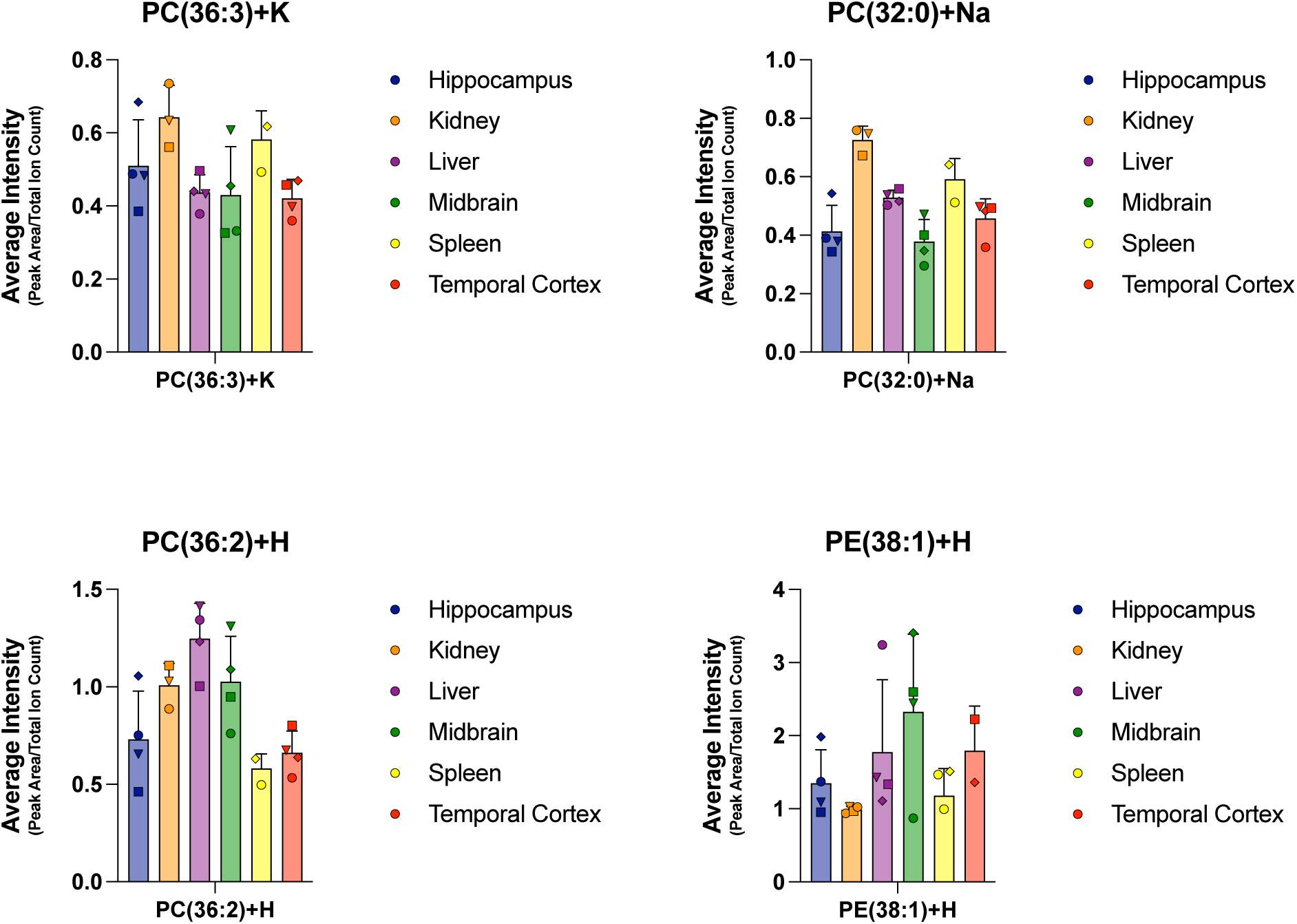
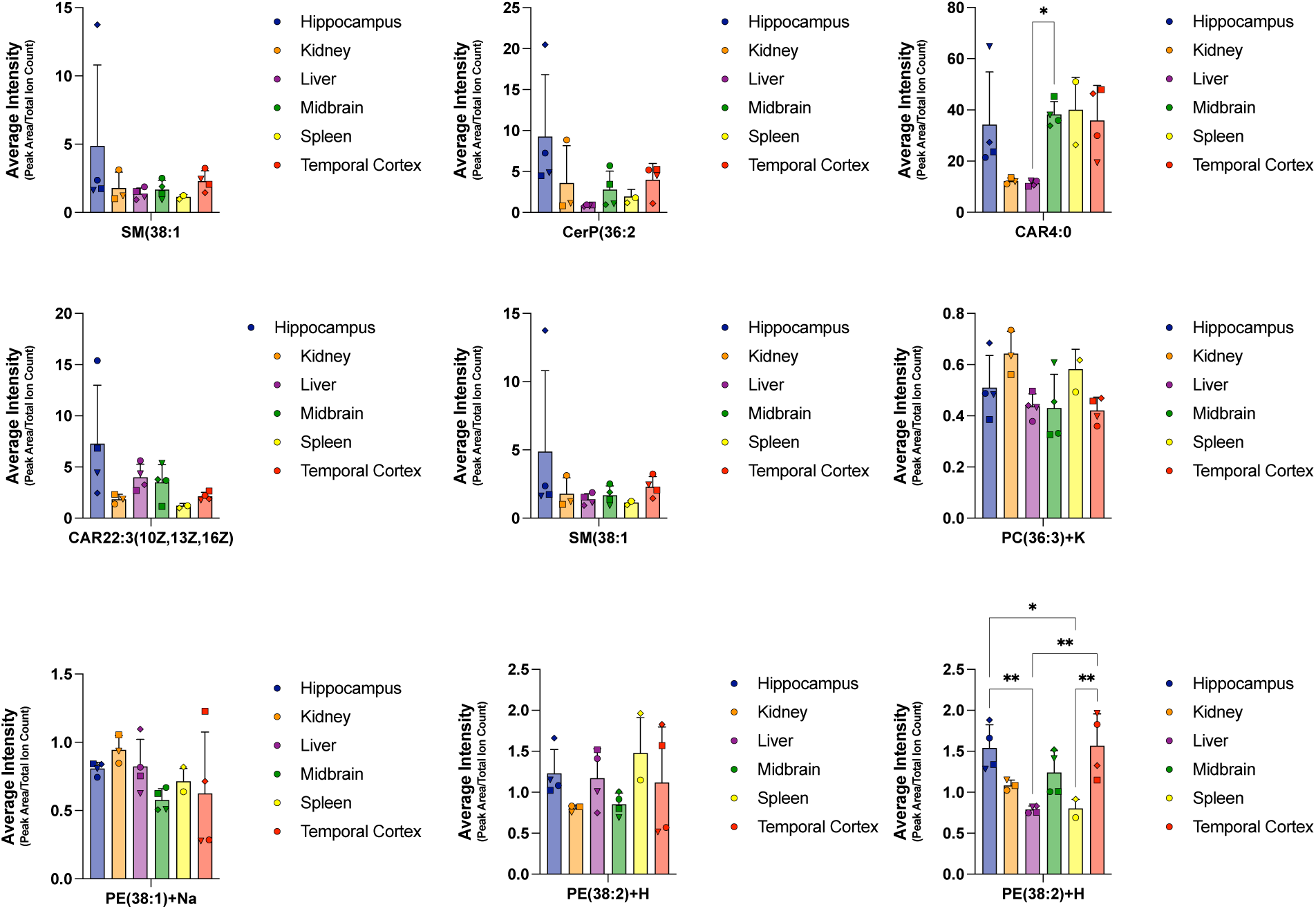
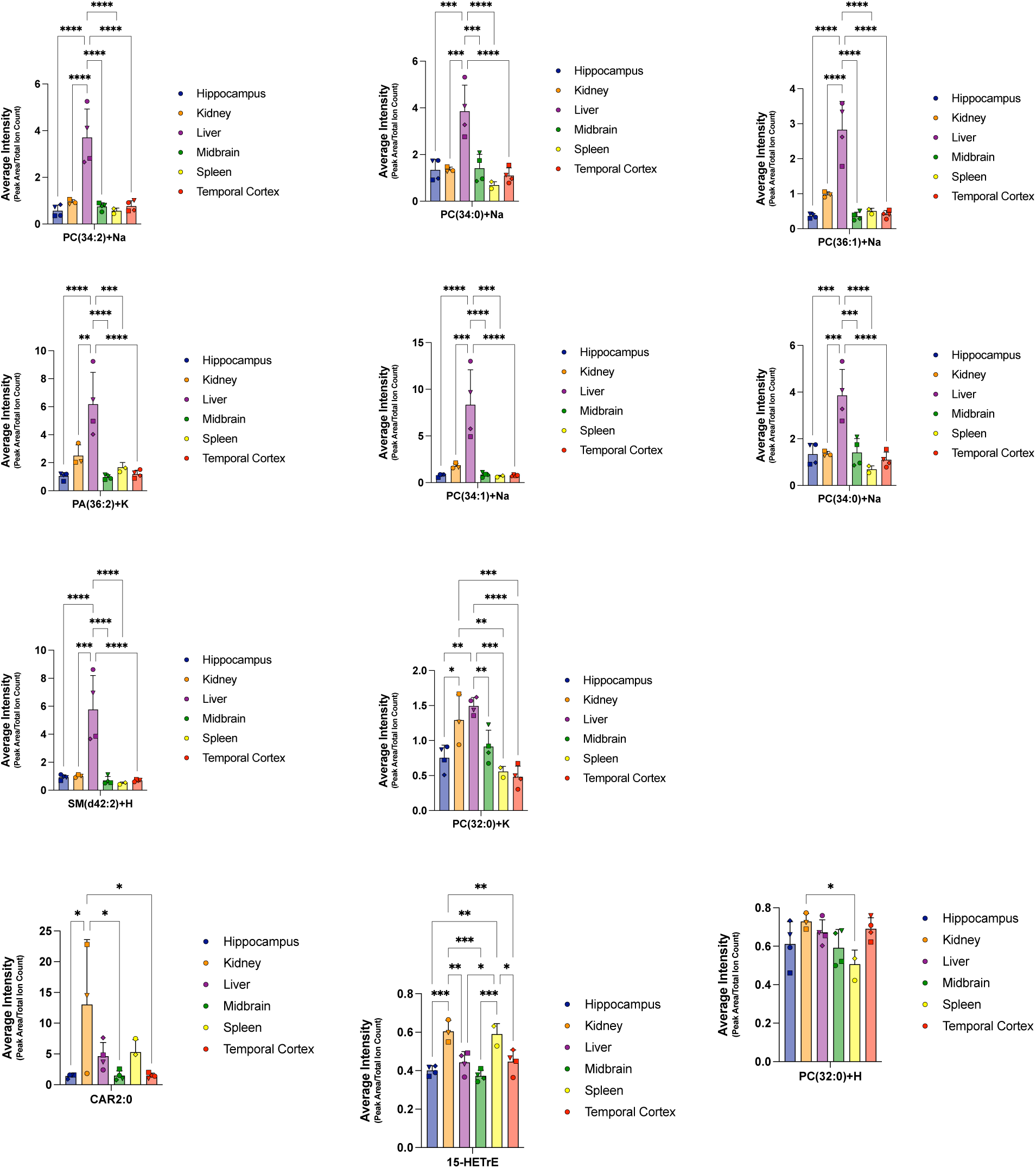
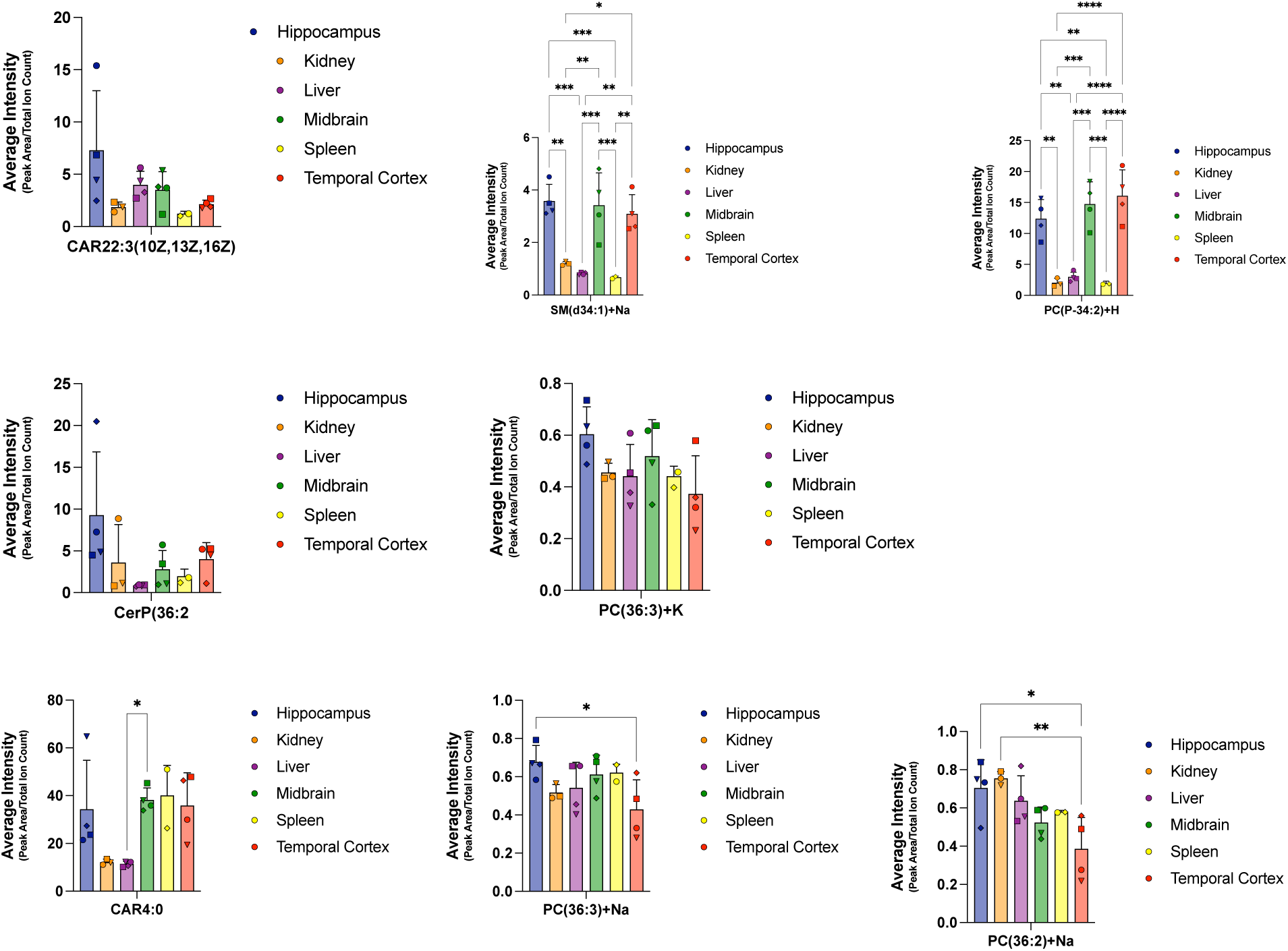
Individual Lipid Intensities Vary by Species, Tissue, and Treatment. Peak areas of lipids identified manually through LipidMaps were used assess overall intensity of ions corresponding to specific individual lipids. All data were analyzed using ordinary one-way analysis of variance (ANOVA) followed by Tukey’s multiple comparisons test. Data are presented as mean. *α = 0.05; **α = 0.01; ***α = 0.001; ****α = 0.0001; ns, not significant.

**Figure S2.**
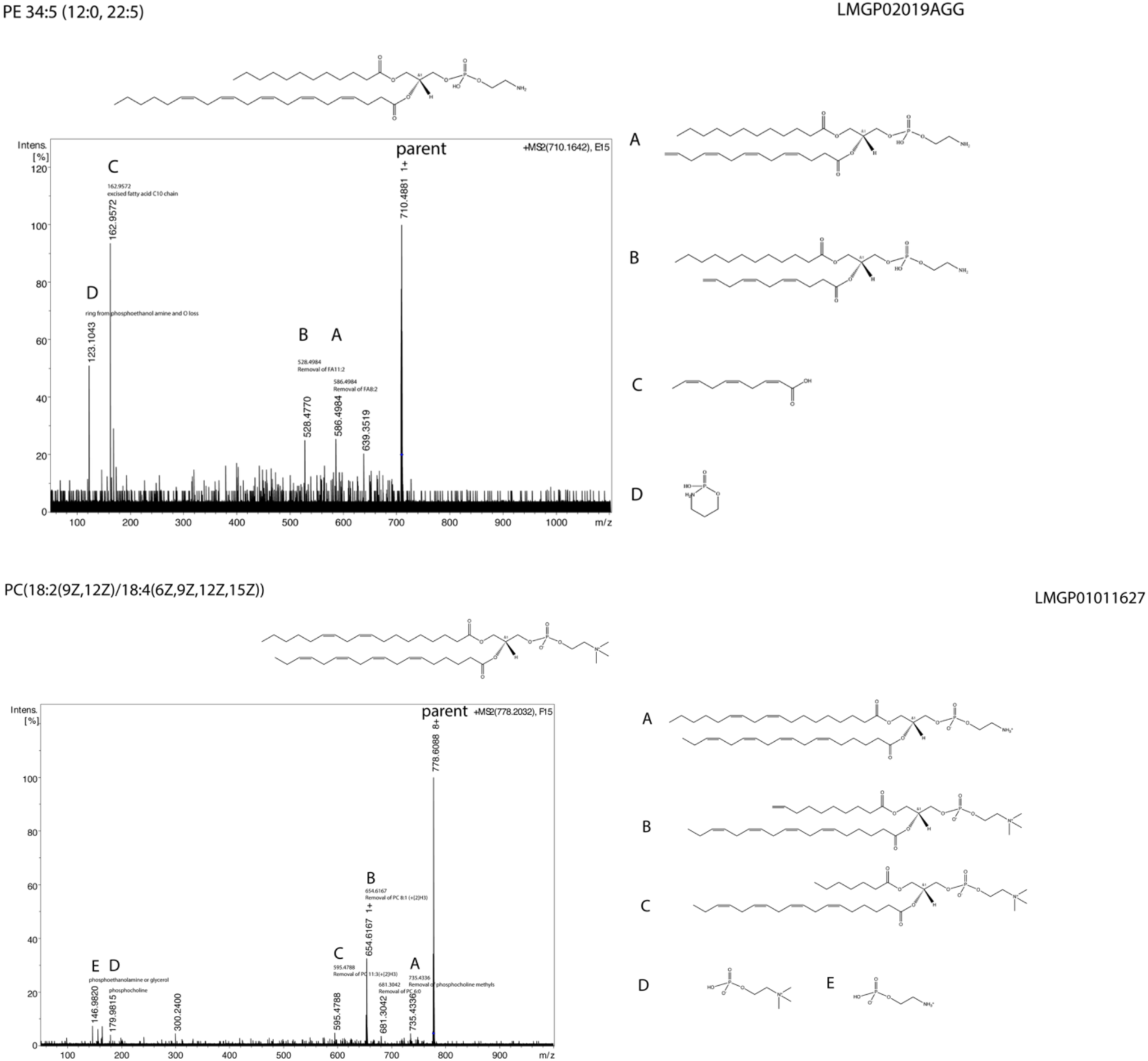
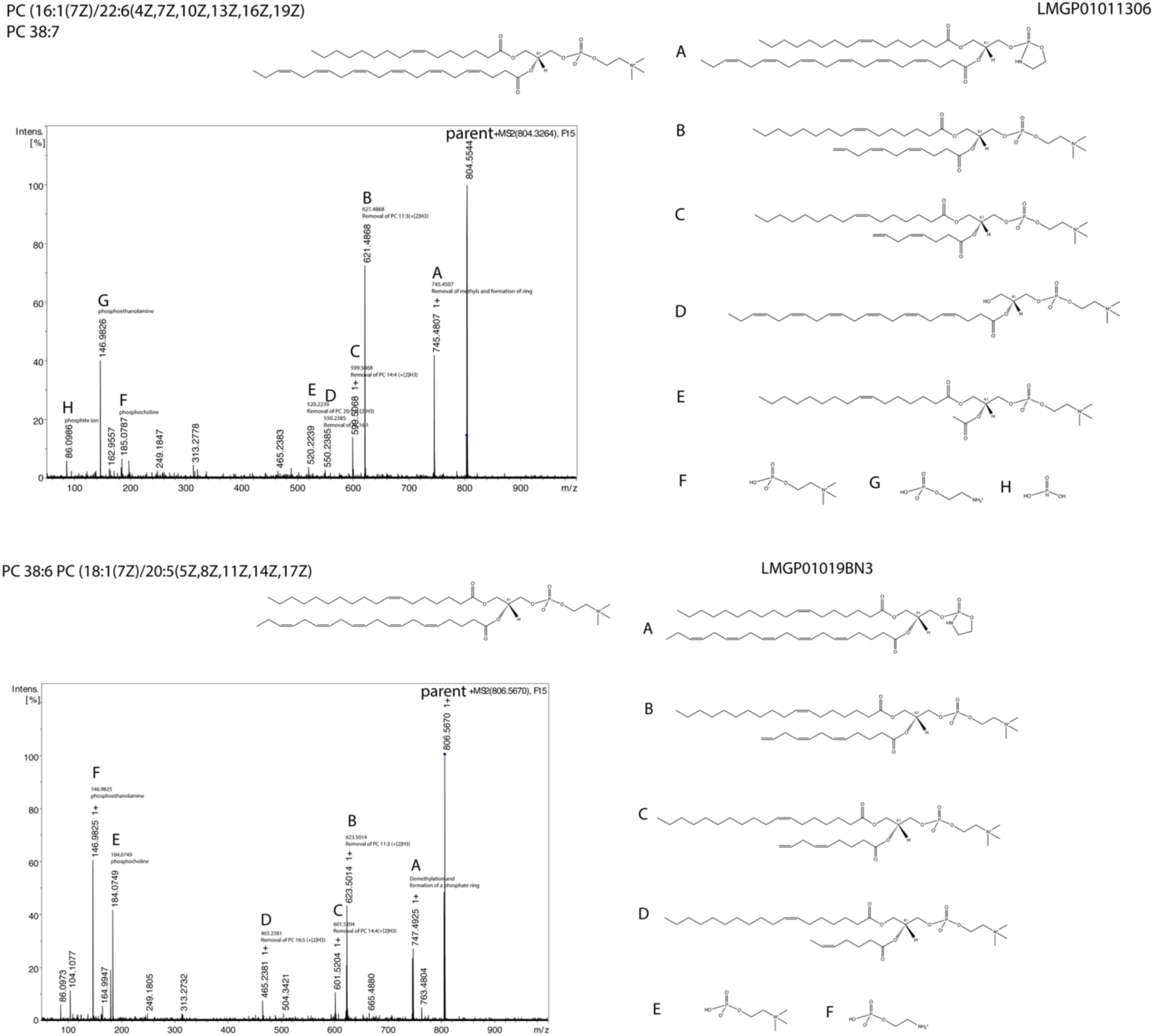
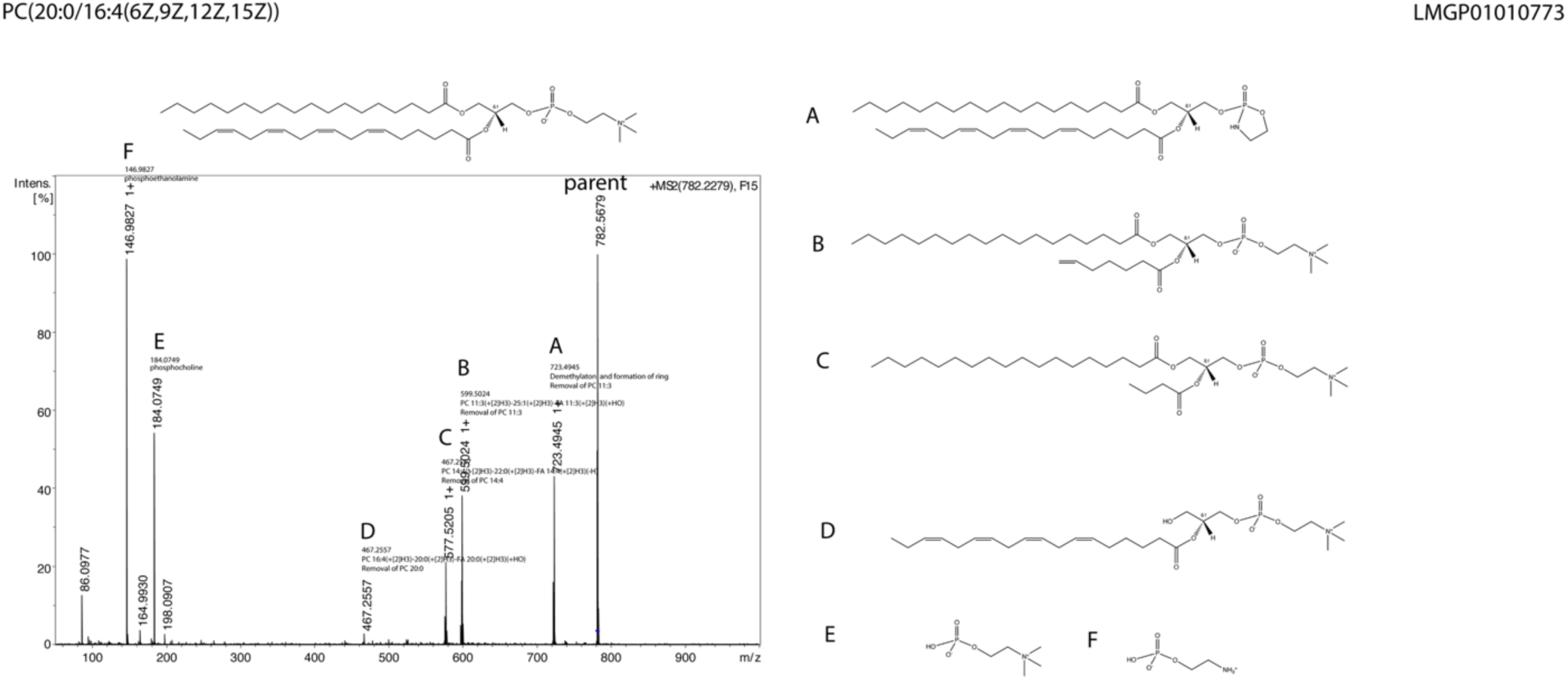
Tandem Mass Spectrometry Validates Identity of Lipids Compared to Corresponding Ions. Labeled chromatograms describe parent lipids manually identified via LipidMaps and corresponding lipid fragments assessed via tandem mass spectrometry (A-H). LipidMap IDs for each respective lipid are listed on each chromatogram.

